# Phylogeny Recapitulates Learning: Self-Optimization of Genetic Code

**DOI:** 10.1101/260877

**Authors:** Oliver Attie, Brian Sulkow, Chong Di, Wei-Gang Qiu

**Author notes:** Emails: Oliver Attie; Brian Sulkow; Chong Di.

## Abstract

Learning algorithms have been proposed as a non-selective mechanism capable of creating complex adaptive systems in life. Evolutionary learning however has not been demonstrated to be a plausible cause for the origin of a specific molecular system. Here we show that genetic codes as optimal as the Standard Genetic Code (SGC) emerge readily by following a molecular analog of the Hebb’s rule (“neurons fire together, wire together”). Specifically, error-minimizing genetic codes are obtained by maximizing the number of physio-chemically similar amino acids assigned to evolutionarily similar codons. Formulating genetic code as a Traveling Salesman Problem (TSP) with amino acids as “cities” and codons as “tour positions” and implemented with a Hopfield neural network, the unsupervised learning algorithm efficiently finds an abundance of genetic codes that are more error-minimizing than SGC. Drawing evidence from molecular phylogenies of contemporary tRNAs and aminoacyl-tRNA synthetases, we show that co-diversification between gene sequences and gene functions, which cumulatively captures functional differences with sequence differences and creates a genomic “memory” of the living environment, provides the biological basis for the Hebbian learning algorithm. Like the Hebb’s rule, the locally acting phylogenetic learning rule, which may simply be stated as increasing phylogenetic divergence for increasing functional difference, could lead to complex and robust life systems. Natural selection, while essential for maintaining gene function, is not necessary to act at system levels. For molecular systems that are self-organizing through phylogenetic learning, the TSP model and its Hopfield network solution offer a promising framework for simulating emerging behavior, forecasting evolutionary trajectories, and designing optimal synthetic systems.

## Introduction

### Limits of natural selection

Discoveries by Alfred Wallace and Charles Darwin in the 19th century established natural selection as the predominant mechanism of evolutionary adaptation (Darwin, 1909; Wallace, 2002). The Modern Synthesis and the more recent Neutral Theory of molecular evolution reaffirm the supremacy of natural selection as the only evolutionary mechanism capable of deterministic, directional changes against a backdrop of stochastic, directionless evolutionary forces including mutation, genetic drift, and recombination (Charlesworth et al., 2017; Huxley, 1974; Kimura, 1984). Yet theories of natural selection and population genetics offer limited explanations on the apparent evolutionary trend towards increasing organizational complexity of living systems, as evident in the repeated emergence of complex and robust structures and molecular processes including molecular pathways, subcellular structures, multicellularity, sexual reproduction, embryonic development, sociality, and self-regulating ecosystems (Pigliucci, 2007; Smith and Szathmary, 1997). A major weakness of Darwinian natural selection and population genetic analysis is its inability to specify testable algorithmic steps, replicate with simulation, or predict the future outcomes of organic complexity (Chaitin, 2013; Valiant, 2013).

In recent years, computational and algorithmic learning emerged as a major contender to bridge the epistemological gap between the feasibility of organismal complexity foretold by the theory of natural selection and its algorithmic plausibility (Chaitin, 2013; Valiant, 2013; Watson and Szathmäry, 2016). Algorithmic learning, defined as the process of creating internal representations (e.g., as memories or genomes) of external regularities through “concrete computation that takes a limited number of steps”, applies equally well to understand the biological origins of cognition and adaptation (Valiant, 2013). By quantifying the degree of computational complexity of the problems imposed by environments, computational learning has not only the potential to generate complex adaptations but also to predict the limit and adverse consequences of adaptations. For example, the learning theory predicts an inability of organisms to adapt to infrequent events (under-fitting) and, conversely, a propensity for organisms to be trapped in mal-adaptions due to biased environments (over-fitting) (Valiant, 2013).

### Evolutionary connectionism

Computational learning is at the heart of a new evolutionary paradigm termed “evolutionary connectionism”, which declares a theoretical equivalence between learning algorithms and evolutionary processes leading to the emergence of complex adaptations (Watson et al., 2016; Watson and Szathmâry, 2016). Central to evolutionary connectionism is the concept of correlational or associative learning first proposed by Donald Hebb in understanding the spontaneous origin of neural networks capable for memory (Hebb, 2002). The Hebbian rule, known colloquially as “cells that fire together, wire together”, is now postulated as a general pattern-generating mechanism in living systems beyond neural systems. For example, the origin of developmental pathways and other gene regulatory networks may be a consequence of following genomic analogs of the Hebbian rule that “genes that are selected together wire together” and “genes that function together junction together” (Vey, 2013; Watson et al., 2014). Bacterial operons, consisting of tandemly arranged and co-transcribed genes, may be a physical manifestation of the Hebbian rule in shaping genome structure (“genes that function together locate together”).

Despite (or perhaps because of) its simplicity, the Hebbian correlational learning has considerable power in generating adaptive and robust features in living systems beyond neural networks. First, as an unsupervised learning algorithm, Hebbian learning is efficient in finding locally optimal solutions without the need to search through a prohibitively large number of possible solutions and test these solutions individually, as is necessary in the process of natural selection. Second, as a distributed algorithm, Hebbian learning occurs locally between directly interacting entities (neurons or genes) while nonetheless leading to emergence of system-level optimal structures (neural or gene networks). It is not necessary to hypothesize that the system-level phenotypes (e.g., developmental processes or gene networks) are themselves direct targets of natural selection. Third, by discovering and encoding trait correlations, Hebbian learning results in networks with foresightedness and robustness by ignoring short-term fluctuations in the optimal values of individual traits (Watson et al., 2016). In short, natural selection operates orthogonally with the principles of self-organization and self-learning in creating complex adaptive systems with coevolving components (Kauffman, 1993; Watson et al., 2016).

For the time being, however, evolutionary connectionism has primarily been a conceptual framework (Vey, 2013; Watson et al., 2014). It remains unclear how extensively Hebbian learning -relative to natural selection and other learning algorithms - operates in shaping molecular and cellular structures beyond neural systems. It is desirable and indeed necessary to test claims of evolutionary learning in the context of a specific molecular system with computational, statistical and biological evidence.

### Origin of genetic code

Here we investigate relative contributions of natural selection and self-learning to the origin of genetic code. Specifically, we test the hypothesis that the standard genetic code may have been evolved predominantly through a Hebbian-learning rather than a natural-selective process. The standard genetic code (SGC, Figure 1A), with which 64 possible combinations of triple nucleotides (“codons”) in DNA encode 20 canonical amino acids in proteins and the translational stop signal, underlines much of the robustness of cells against deleterious mutations. For example, single-nucleotide mutations occurring at the 3^rd^ codon position typically result in no change of amino acids. Such properties of SGC are called “error-minimization”, referring to the fact that SGC is non-randomly structured to minimize disruptions by DNA mutations to protein sequences, structures, and functions.

**Figure 1.**
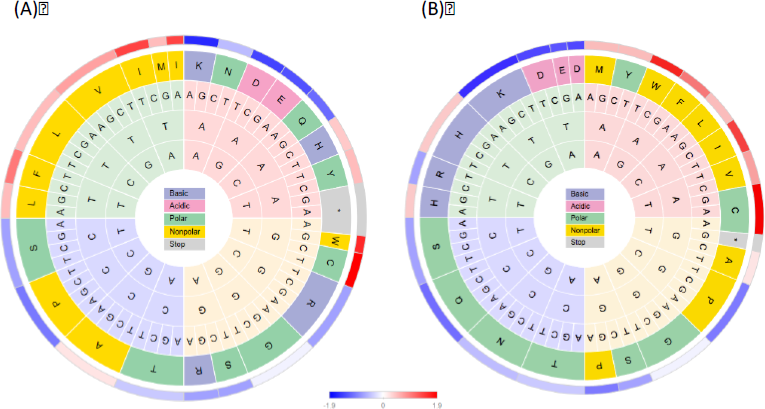
The standard genetic code (SGC) and a Hopfield-optimized genetic code, shown with a codon wheel that minimizes evolutionary distances between codons. **(A)** Standard genetic code. Nucleotide bases are arranged (from center outward) in the order of 1^st^, 2^nd^, and 3^rd^ codon positions. Within each of these three rings, the four bases are repeated clockwise in the order of “AGCT” to minimize the number of transversions, which has a greater evolutionary distance than transitions. Base cycling is the fastest at the 3^rd^ and the slowest at the 2^nd^ codon positions, reflecting the decreasing evolutionary rates in the order of the 3^rd^, the 1^st^, and the 2^nd^ positions. With these rules, the codon wheel represents the shortest mutation path and is analogous to the time dimension in a Traveling Salesman Problem (TSP). At the 4^th^ ring, the 20 amino acids are colored physio-chemically according the legend in the center. At the 5^th^ ring, amino acids are colored by a continuous scale of hydrophobicity indicated in the bottom (Casari and Sippl, 1992). **(B)** An optimal genetic code generated using a Hopfield network that minimizes polarity and volume errors. This simulated code is more optimized than SGC in e.g., displaying more consistent amino acid physio-chemistry for the bases at the 2^nd^ codon position: all “-A-” codons code for hydrophobic amino acids, the majority of “-G-” and “-C-” codons code for hydrophilic amino acids, and all “-T-” codons code for charged amino acids, with “AT-” coding for the two negatively charged amino acids and the rest coding for positively charged amino acids. In SGC, codons encoding the three positively charged amino acids (H, K, and R) are separated by large mutation distances.

While there is no question on the adaptiveness of SGC, the molecular processes that led to its origin remain controversial (Freeland et al., 2000; Goldenfeld et al., 2017; Koonin, 2017). Natural selection, in conjunction with chemical processes during early history of life, has been the most frequently argued hypothesis to explain the origin of the highly optimized and nearly universal SGC (Di Giulio, 2016; Freeland et al., 2000; Koonin, 2017). However, the selective mechanism tends to be slow and inefficient since it implies that SGC emerges from competition among cells equipped with random genetic codes (Massey, 2016). More contentiously, since the protein translation machinery is a complex subcellular system consisting of, among others, tRNAs, aminoacyl-tRNA synthetases (aaRSs) and the ribosome, the selective hypothesis raised the question whether (and how) natural selection operates at the level of individual genes, the subcellular system, or the cell system as a whole (Kauffman, 1993). For example, it has been argued that the apparent adaptiveness of SGC may have risen as a byproduct of incremental evolution through codon capture, during which structurally similar aaRSs tend to recognize phylogenetically similar tRNAs and physio-chemically similar amino acids (Davis, 1999; Osawa and Jukes, 1989; Stoltzfus and Yampolsky, 2007). Regardless, neither the selective hypothesis nor the incremental evolution hypothesis directly address the algorithmic processes by which error-minimizing capacity of SGC has evolved. Besides being of considerable evolutionary interest, the design principle and algorithmic origin of SGC are of practical importance for reengineering the genetic code to create synthetic therapeutic proteins using non-canonical amino acids, among other applications (Chin, 2017; Xue and Wong, 2017).

Here, we explore the possibility of an evolutionary origin of SGC through Hebbian learning. Specifically, we propose that Hebbian learning captures the algorithmic essence of the codon capture and reassignment hypotheses, both of which imply an incremental phylogenetic process of evolving aaRSs and tRNAs recognizing increasingly dis-similar amino acids (Davis, 1999; Osawa and Jukes, 1989; Stoltzfus and Yampolsky, 2007). The search for optimal genetic codes by using learning algorithms is not new and these studies generally conclude that SGC is far from being globally or even locally optimal (de Oliveira et al., 2015; Santos and Monteagudo, 2017; Tlusty, 2010). Our primary objective is not to propose a new learning algorithm or to find more optimal genetic codes but to demonstrate that Hebbian learning is a key design principle that explains both the evolutionary origin of SGC and its error-minimizing robustness.

## Models & Methods

### Hopfield network & Traveling Salesman Problem

Hopfield neural network is an algorithmic model of associative memory that implements the Hebbian learning rule (Hopfield, 1982). A Hopfield neural network, constructed (electronically or computationally) with a set of symmetrically connected neurons, is capable of storing, retrieving, and restoring memories when activities of individual neurons are determined by the Hebbian learning rule (Hopfield, 1982; MacKay, 2003). For example, in a binary Hopfield network where each neuron’s activity takes the value of either 1 or −1, the connection weight (*w_i,j_*) between a pair of neurons is increased if their activities (*x_i_* and *x_j_*) are positively correlated (both 1 or both −1) and decreased if negatively correlated (one 1 and the other −1). Once connection weights are determined with a set of stored memories (vectors of 1’s and −1’s), from a randomly set initial state the network is mathematically guaranteed to reach a locally stable state where the following definition of the network energy is at the minimum (MacKay, 2003):

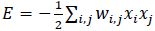

As such, while the neuron activities are determined at local levels through interactions with each other, the neural network displays collective, system-level behavior mimicking associative memory. The network is able to, for example, retrieve a complete state from incomplete inputs or recover a correct state from mistaken inputs. More surprisingly and perhaps more importantly, the system is robust in the sense that the network functions well even when some of the neurons are removed, mimicking a damaged yet functional brain and indicating information redundancy encoded in the network (MacKay, 2003).

Besides being a powerful model of associative memory, Hopfield network is an efficient algorithm for solving combinatorial optimization problems, such as the traveling salesman problem (TSP), which is to find the shortest tours to cover *N* cities while visiting each city once (Hopfield and Tank, 1986). Instead of being used as a memory device, here the Hopfield network is used as a computational tool to search for optimal multivariate states. To solve the TSP, for example, a Hopfield network is simulated with *N^2^* neurons, each representing the probability of a city being visited at a tour position. An energy function is defined to reflect both the constraints (e.g., each city to be visited once and the salesman can visit only one city at a time) and a measurement to be minimized (e.g., the tour length). When initialized with arbitrary activities, the network progressively reaches a minimum representing a locally shortest path as the energy function converges to a stable local minimum (Hopfield and Tank, 1986; Potvin, 1993).

The original Hopfield-Tank algorithm was found difficult to replicate as the number of cities increases and a more efficient algorithm for finding shortest paths using neural normalization and simulated annealing was proposed (Bout and Miller, 1989). The mean field of a neuron representing city *X* visited at tour position *i* is defined as:

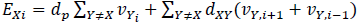
where *d_p_* is a penalty experimentally adjusted to ensure that only one city can occupy a tour position and *v_xi_* is the probability of city *X* at tour position *i*. Values of *v_xi_*’s are assumed to obey a Boltzman distribution at any given simulated temperature T: *v_xi_ α e^−E_xi_/T^*. At each iterative updating step, the neuron outputs are normalized to sum up to one so that each value represents a true probability:

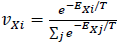

Finding a valid path depends on the temperature *T* and the penalty dp. As the temperature increases, neuron outputs become increasingly uniform (*v_xi_* 1/n). As the temperature drops to a critical value (*T_0_*), the neurons anneal to a steady low-energy state representing a stable, locally minimal-energy mapping between the cities and tour positions. Both the critical temperature *T_0_* and *d_p_* that lead to valid tours are experimentally determined by simulations (Bout and Miller, 1989).

### Algorithm to generate optimal genetic codes using TSP

#### TSP model of SGC origin

The genetic code could be modeled as a tour in the Traveling Salesman Problem (TSP) so that it could be optimized by using a Hopfield neural network. We hypothesize that an optimal genetic code minimizes the total distance of a tour of 20 amino acids from one codon to next, analogous to a tour of *N* cities by a salesman from one time point to the next. To make the codons equivalent to the linearly ordered time points in a TSP, it is necessary to generate a linearly ordered sequence of codons that minimizes the cumulative mutational distance between codons (see “codon wheel” below).

To represent genetic codes as a TSP, we first construct a Hopfield network consisting of 21 x 21 neurons, the activity of each of which representing the probability of an amino acid (“cities”) at a tour position. We measured distances between two amino acids (X and Y) by the Euclidean distance: *d_XY_* = 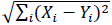, where *i* stands for one of the physio-chemical indices (either hydrophobicity, polarity, volume, or isoelectric point; Table 1). To remove effects of difference in magnitudes when combining multiple indices, we rescale each index by normalizing the values to a mean of zero while maintaining the standard deviation. Values for stop codons are arbitrarily assigned to be an outlier (greater or less than two standard deviations from the mean).

**Table 1.** Physio-chemical indices of amino acids^a^.

| 1-letter code | 3-letter code | Polarity | Hydrophobicity | Volume | Iso-electric point |
| --- | --- | --- | --- | --- | --- |
| A | Ala | 7.0 (-0.1672) | 1.8 (0.7638) | 31 (-1.067) | 6.00 (-0.02972) |
| C | Cys | 4.8 (-1.043) | 2.5 (1.002) | 55 (-0.5477) | 5.07 (-0.5561) |
| D | Asp | 13.0 (2.221) | -3.5 (-1.039) | 54 (-0.5694) | 2.77 (-1.858) |
| E | Glu | 12.5 (2.022) | -3.5 (-1.039) | 8.3 (-1.559) | 3.22 (-1.603) |
| F | Phe | 5.0 (-0.9631) | 2.8 (1.104) | 132 (1.120) | 5.48 (-0.3241) |
| G | Gly | 7.9 (0.1910) | -0.4 (0.01531) | 3 (-1.674) | 5.97 (-0.0467) |
| H | His | 8.4 (0.3900) | -3.2 (-0.9373) | 96 (0.3402) | 7.59 (0.8703) |
| I | Ile | 4.9 (-1.003) | 4.5 (1.682) | 111 (0.6651) | 6.02 (-0.0184) |
| K | Lys | 10.1 (1.067) | -3.0 (-0.8693) | 119 (0.8384) | 9.74 (2.087) |
| L | Leu | 4.9 (-1.027) | 3.8 (1.444) | 111 (0.6651) | 5.98 (-0.04104) |
| M | Met | 5.3 (-0.8437) | 1.9 (0.7978) | 105 (0.5352) | 5.74 (-0.1769) |
| N | Asn | 10.0 (1.027) | -3.5 (-1.039) | 56 (-0.5261) | 5.41 (-0.3637) |
| P | Pro | 6.6 (-0.3264) | -1.6 (-0.3930) | 32.5 (-1.035) | 6.30 (0.1401) |
| Q | Gln | 8.6 (0.4696) | -3.5 (-1.039) | 85 (0.102) | 5.65 (-0.2278) |
| R | Arg | 9.1 (0.6686) | -4.5 (-1.380) | 124 (0.9466) | 10.76 (2.665) |
| S | Ser | 7.5 (0.03184) | -0.8 (-0.1208) | 32 (-1.046) | 5.68 (-0.2108) |
| T | Thr | 6.6 (-0.3264) | -0.7 (-0.08676) | 61 (-0.4178) | 6.16 (0.06085) |
| V | Val | 5.6 (-0.7243) | 4.2 (1.580) | 84 (0.08035) | 5.96 (-0.05236) |
| W | Trp | 5.2 (-0.8835) | -0.9 (0.1548) | 170 (1.943) | 5.89 (-0.09198) |
| X<sup>b</sup> | Stop | (-1.5993) | (1.672) | (-1.5833) | (-2.6562) |
| Y | Tyr | 5.4 (-0.8039) | -1.3 (0.2909) | 136 (1.207) | 5.66 (-0.2222) |

<sup>a</sup> Raw values were obtained from (Haig and Hurst, 1991) and normally scaled indices are in parenthesis. Stop signal was excluded from normalization. These indices are not independent from each other, as shown by a principal component analysis (not shown).
<sup>b</sup> Stop signal, assigned values that are statistical outliers (least polar, most hydrophobic, largest in volume, and most negatively charged) so that its distances from amino acids are greater than the distance between any two amino acids.

Second, the neural network is initialized with uniformly distributed random activities centered at 0.5 (Hopfield and Tank, 1986). The network is then optimized by following the simulated annealing algorithm with preset values of *d_p_* (e.g., dp=0.7) and T(e.g., T=0.1), determined experimentally to maximize the proportion of valid tour paths (Equations 2 & 3) (Bout and Miller, 1989). Each resulting optimal tour path is checked for validity to ensure that all amino acids are covered and each amino acid is visited only once. Invalid paths are discarded and valid paths are saved for further processing.

Third, we map the amino acids from an optimal path produced by the Hopfield network to the 64 possible codons. Since there are more codons than amino acids it is necessary to assign multiple codons to a single amino acid.

#### Codon wheel

We construct a linear order of codons with minimal evolutionary distances by following known molecular evolutionary principles. First we give preference to mutations at the 3^rd^ codon position and then to those at the 1^st^ codon position, followed by those at the 2^nd^ codon position, reflecting increasing evolutionary distances of nucleotide substitutions from the 3^rd^, to the 1^st^, and the 2^nd^ codon positions. Second, for mutations introduced to the same codon positions, we give preference to transitions over tranversions, reflecting smaller evolutionary distances of transitions than transversions. Following these two rules, a linear sequence of codons, cycling the four bases at each codon position by the order of “AGCTTCGA” - a way to minimize the number of transversions - are uniquely defined and shown as a circular codon wheel (Figure 1). The codon wheel could alternatively be defined by any of the other three possible transversion-minimizing base-cycling orders (“AGTCCTGA”, “GACTTCAG”, and “GATCCTAG”).

To assign multiple codons to the same amino acid, we start with an arbitrary codon in a codon wheel (e.g., “AAA”) and travel clockwise through all codons, while labeling each codon with an integer determined by the order of distinct amino acid this codon is assigned to according to SGC (Figure 1). In other words, we assign each codon an “SGC address”, which may or may not be shared with another codon. For example, starting from “AAA” in a codon wheel shown as Figure 1, the codons are labeled as “AAA” (1), “AAG” (1), “AAC” (2), “AAT” (2), “GAT” (3), “GAC” (3), “GAG” (4), “GAA” (4), and so forth and end with “ATT” (19), “ATC” (19), “ATG” (20), and “ATA” (21). As such, an optimal amino-acid path generated by Hopfield network could be assigned to 64 codons based on tour positions. For example, if Lysine (K) has a tour position of 4 in a TSP path, it will be assigned to two codons “GAG” and “GAA”, both of which have an “SGC address” of 4, although Lysine (K) is assigned to “AAA” and “AAG” in SGC. Note that the SGC addresses of codons are arbitrarily assigned as long as they follow the order of a codon wheel. If tour positions of amino acids are random (i.e., not optimized by Hopfield network), this scheme is equivalent to a random permutation of amino acids among synonymous codon blocks (Freeland et al., 2000). A Hopfield-optimized tour path of amino acids, on the other hand, is expected to be ordered to maximize the chance that similar amino acids are assigned to similar codons (Stoltzfus and Yampolsky, 2007).

### Statistical analysis of genetic codes

#### Randomized genetic codes

To test optimality of SGC and simulated genetic codes, we generate random codes as statistical controls by permuting the 20 amino acids and the stop signal among the 21 synonymous codon blocks (Freeland et al., 2000). This randomization scheme is a stringent test of code optimality. It maintains the same codon degeneracy as in SGC while disrupting any correlation that might exist between amino acids and codon blocks (Freeland et al., 2000).

#### Code optimality measured by mutational error

We quantify optimality of each code by calculating errors (i.e., changes in an index value) caused by single-nucleotide substitutions (Haig and Hurst, 1991). The mutational error of a code, a measure of overall code fitness, is the average error across all pairs of single-mutation codons (Freeland et al., 2000):

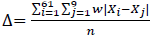
where *i* is the source codon, *j* is the destination codon (stop codons excluded) differing from the source codon by a single nucleotide, w is the ti/tv bias, *X_i_* and *X_j_* are the physio-chemical values of the two amino acids associated with the two codons, and *n* is the total number of one-nucleotide neighboring codon pairs. Statistical significance of the error of a code is assessed by the proportion of random codes with an equal or smaller error. Errors are also be calculated individually at the three codon positions and for transitions and transversions separately as a way to estimate if errors are minimized at individual codon positions and for transition or transversion.

### Phylogenetic analysis

To explore biological basis of evolutionary learning, we infer early evolutionary events during the origin of SGC using molecular sequences of contemporary aminoacyl-tRNA synthetases (aaRSs) and tRNAs from the model bacterial species *Escherichia coli.* We download aaRS sequences from EcoGene (http://www.ecogene.org/), the online database of *E.coli* K-12 gene sequences (Zhou and Rudd, 2013). The aaRSs are subdivided into two classes according to a structure-based classification (Nagel and Doolittle, 1995). Protein sequences within each aaRS class are multiply aligned with MUSCLE (Edgar, 2004) and an approximate maximum likelihood phylogenetic tree is obtained with FastTree (Price et al., 2010) using default settings. Using the BpWrapper BIOTREE utility (Hernandez et al., 2018), branches with low bootstrap support (<0.7) are collapsed and the tree is rooted at the midpoint of between two most-distant nodes.

We download structurally aligned tRNA gene sequences from the same species and strain (*E.coli* K-12) from tRNAdb (Jühling et al., 2009). Redundant sequences are removed and an approximate maximum likelihood phylogenetic tree is obtained with FastTree (Price et al., 2010) using default settings. Using the BpWrapper BIOTREE utility (Hernandez et al., 2018), branches with low bootstrap support (<0.7) are collapsed and the tree is rooted at the midpoint. All phylogenetic trees were plotted using the APE package on the R and RStudio platform (Paradis, 2012).

We use phylogenetic autocorrelation to test co-diversification of aaRS and tRNA sequences with their cognate amino acids. Phylogenetic autocorrelation is a measure of association of a variable with a phylogeny, in the same way as spatial autocorrelation measures influence of a variable by geographic distances (Gittleman and Kot, 1990; Moran, 1950). The same statistics is employed for both purposes with one based on geographic distances and the other on tree distances. We use the *gearymoran* function in ADE4 to calculate Moran’s I with an amino acid index (e.g., polarity) and obtained its statistical significance with Geary’s randomization protocol (Dray and Dufour, 2007).

### Results

### Two sources of error minimization in SGC

There are a total of 263 pairs of amino-acid encoding codons that differ by a single base. These codon pairs could be categorized into six types according to the position of the differing base and whether it is a transition (A/G or C/T) or transversion (A/C, A/T, G/C, or G/T). Transitions occur more frequently than transversions, known as the ti/tv mutational bias. When the error magnitude (measured by e.g., amino-acid polarity; Table 1) between two codons in SGC are plotted for each of the six categories, it is apparent that mutations at the 3^rd^ codon position cause the least errors, followed by the 1^st^ codon position and then by the 2^nd^ codon position (red boxes in Figure 2A). This is largely due to codon degeneracy by which multiple codons code for the same amino acid. For example, 29 the 30 possible single-nucleotide transitions and 34 out of 58 possible transversions at the 3^rd^ codon position are synonymous in SGC. At the 1^st^ codon position, two out of 29 transitions (CTA/CTG & CTG/TTG, all encoding Leu) and two out of 58 transversions (AGA/CGA & AGG/CGG, all encoding Arg) are synonymous (Figure 1A). There is no codon degeneracy at the 2^nd^ codon position. Apparently, codon degeneracy in SGC is non-uniform, decreasing in the order of the 3^rd^, 1^st^, and 2^nd^ codon positions and over-represented in transitions. Codon degeneracy, however, is not itself adaptive since some form of degeneracy is inevitable for any genetic code consisting of more codons than amino acids.

**Figure 2.**
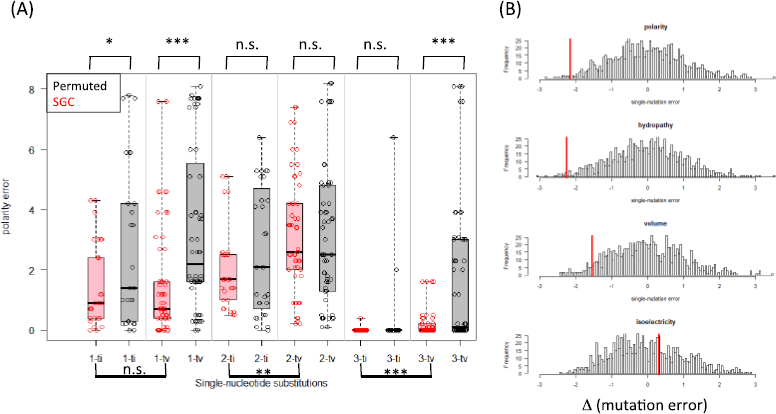
Patterns of mutation-error minimization in SGC. **(A)** On the x-axis, single-nucleotide mutations between pairs of codons are categorized by codon positions (1, 2, and 3) and by transitions (ti) or transversions (tv). The y-axis shows the errors caused by such mutations as quantified by the polarity index (Table 1). Points in red (n=263) are derived from SGC; points in black (n=258) by a single round of random permutation of amino acids among the synonymous codon blocks (Freeland et al., 2000). This randomization scheme maintains the blocks of synonymous codons while disrupting any patterns present between neighboring codon blocks. Two sources of error minimization are identified from this analysis: (i) SGC significantly reduces mutation errors at the 1st codon position and transversion errors at the 3rd codon position (*, 0.01 < p < 0.05 by t-test; **, 0.001 < p < 0.01; p < 0.001), reflecting high levels of code degeneracy at these two positions. (*ii*) At the 2nd and 3rd codon positions, errors by transitions are significantly minimized relative to those by transversions, reflecting an adaptation to a higher rate of transitions relative to transversion in DNA mutations. **(B)** Distributions of amino-acid errors caused by single-nucleotide substitutions (calculated by EQ4 with ti/tv=5) from 1000 permuted codes. Red vertical lines indicate error values for SGC. These error distributions show SGC significantly reduces mutation errors for polarity (p=0.010), hydrophobicity (p=0.011), and volume (p=0.048), but not for isoelectric points (p=0.648).

If SGC does not minimize errors between neighboring codon blocks, one would expect the mutational errors similar between SGC and a randomized code. In reality, errors in SGC (measured by, e.g., errors in polarity) are significantly reduced relative to the permuted codes for transitions and transversions at the 1^st^ codon position (p=2.9e-2 and 8.7e-5 by t-tests) and transversions at the 3^rd^ codon position (p=1.4e-5), while there is no significant error reduction at the 2^nd^ codon position (Figure 2A). For example, at the 3^rd^ codon position, the only non-synonymous transition (ATA[Ile]/ATG[Met]) results in only a minor shift in amino acid polarity (by 0.157 standard deviation; Table 1). This pattern of error reduction indicates that SGC minimizes errors caused by mutations at the 1^st^ and 3^rd^ codon positions in addition to codon degeneracy at these positions. At the 2^nd^ codon position, although there is neither codon degeneracy nor significant error reduction relative to the random code, transitions cause significantly less errors than transversions (p=1.3e-3).

In sum, two sources of error minimization in SGC are identified from this analysis: (*i*) SGC significantly reduces mutation errors at the 1^st^ codon position and transversion errors at the 3^rd^ codon position and (*ii*) at the 2^nd^ and the 3^rd^ codon positions, errors by transitions are significantly minimized relative to those by transversions. These two rules are used to construct a codon wheel that minimizes evolutionary distances between codons (Figure 1).

### SGC co-minimizes errors in amino acid polarity, hydrophobicity, and volume

We test the overall errors of SGC relative to random codes for the four major amino acid indices (Table 1). Results indicate that SGC significantly minimizes errors in polarity (p=0.010), hydrophobicity (p=0.011), and volume (p=0.048), but not iso-electricity (p=0.648) (Figure 2B). The co-minimization of polarity and hydrophobicity is not surprising since these two indices are significantly anti-correlated, according to a principle component analysis of the four indices (not shown).

### Hopfield-optimized genetic codes

We use Hopfield networks to search for optimal genetic codes with the goal of assessing the possibility that SGC may have originated by a similar self-learning process. First, we obtain optimal codes by minimizing the total distance measured by a single parameter. For example, a total of 856 valid paths are generated from a run of 2500 rounds of simulations using amino acid distances determined by polarity. Indeed, the polarity errors of the simulated codes decrease significantly relative to the random codes and the mean error rate is close to that of SGC (−2.0 standard deviation from the mean error of random codes) (Figure 3A). The errors for hydrophobicity, volume, and iso-electricity of simulated codes are incidentally reduced. Adding volume as an extra distance parameter has similar error-reducing efforts to all four indices while the decrease in volume is the largest (−3.0 standard deviation from the mean error) (Figure 3B). Continue by adding iso-electricity as the 3^rd^ parameter, the simulated codes improve upon the random codes for all three measures of errors (Figure 3C). Finally, including all four parameters results in simulated codes optimized for all parameters (Figure 3D).

**Figure 3.**
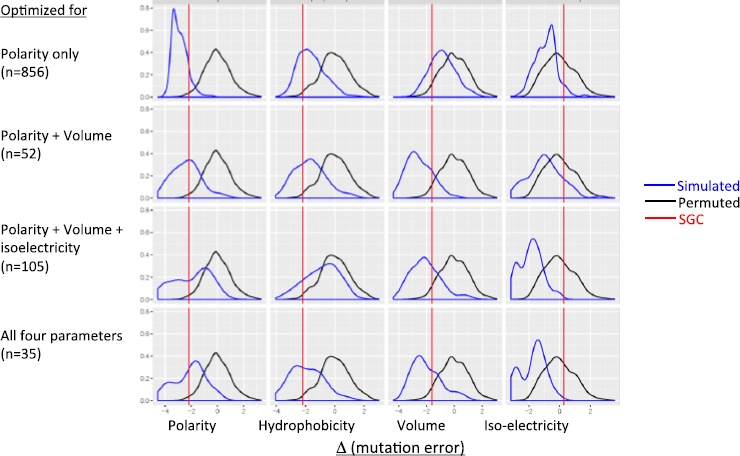
Hopfield network minimizes code errors for any given amino acid indices. We constructed a 21-by-21 TSP and searched for the shortest paths traversing the 20 amino acids and the stop codon (21 “cities”) using Hopfield neural networks (see Methods). Distances between the “cities” were determined by either polarity alone (top row), or a combination of 2, 3, and all 4 parameters (other rows). Valid paths (number in parenthesis) were mapped to the codon wheel resulting in simulated genetic codes. Each simulated code was calculated for average single-mutation error rates based on each parameters (EQ 4). All error rates were normalized according to the mean and standard deviation of random codes (as in Figure 2B) and their distribution were shown as density plots. The Hopfield network is highly responsive to changes in distance function, as seen by a decrease in errors in the parameter(s) included in the distance function. Since these parameters are not entirely independent (e.g., polarity and hydrophobicity are anti-correlated), errors in some excluded parameters were incidentally reduced. While it is possible to obtain codes optimized for all four parameters (bottom row), the most parsimonious codes with errors similar to SGC errors are generated with the Hopfield network that optimizes for polarity alone (top row).

It could be concluded from these simulations that Hopfield network optimizes genetic code in a highly responsive manner depending on which and how many indices are included in the distance function. Further, considering the large iso-electricity error rate of SGC (0.3 standard deviation greater than the random error, Figure 2B), we reject the hypothesis that SGC evolved by optimizing errors in isoelectric point. Indeed, it is most parsimonious to conclude that SGC evolved by minimizing the polarity error alone, with reduced errors in hydrophobicity and volume as incidental consequences (Figure 3A). This hypothesis is supported by the fact that optimization with respect to hydrophobicity or volume alone result in poorer match of errors between SGC and the simulated codes (not shown).

Next, we search for simulated codes that are more optimal than SGC by plotting the error rates grouped by individual codes (Figure 4). One such code with all four errors less than −2.0 standard deviation away from the mean random errors is identified and its codon assignment is visualized with a codon wheel (Figure 1B). This Hopfield-optimized code is more optimal than SGC in that, e.g., all non-polar amino acids are mapped to codons with the 2^nd^ codon position being a purine (A or G). Furthermore, all positively or negatively charged amino acids are mapped to codons with the 2^nd^ codon position being a thymine (T). Most significantly, genetic codes similarly or more optimal than SGC are not a rarity but emerge readily from a Hopfield network (Figure 4), suggesting that SGC is likely suboptimal in error minimization.

**Figure 4.**
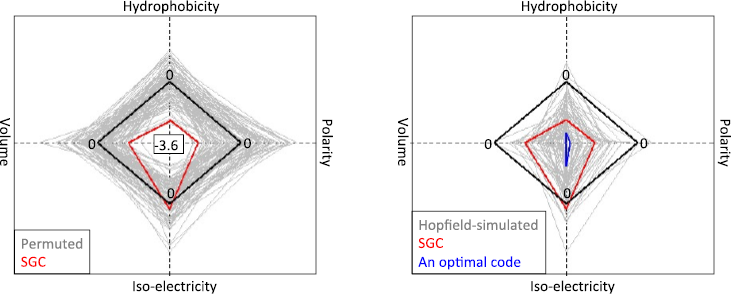
Hopfield network generates genetic codes more optimal than SGC. **(Left Panel)** In this alternative display of error rates of genetic codes, each code is represented by a quadrilateral with vertices defined by the four errors. The black diamond represents the average errors of the permutated codes (n=100). Note that the minimum error (−3.6 standard deviation from the mean) is placed at the center for all four parameters so that the smaller a quadrilateral condenses towards the center the more optimal a code is. **(Right Panel)** Gray quadrilaterals represent Hopfield-simulated codes (n=52) optimized for polarity and volume (the same output used for Figure 3, 2^nd^ row). The simulated codes have generally less errors than randomized codes. Many are more optimal than SGC, one of which is highlighted in blue and its coding pattern is shown with a codon wheel in Figure 1B.

### Phylogenetic autocorrelations between macromolecules and their cognate amino acids

So far we have shown that SGC is an unsurprising (and indeed suboptimal) outcome of a Hebbian learning process when operating under the same set of biological constraints as in SGC. Here we explore possible biological basis of Hebbian learning during the origin of SGC by examining footprints of early evolutionary events left in the gene sequences of the contemporary proteintranslation system from a single *E.coli* genome.

#### aaRSs Phylogeny

A salient feature of Hebbian and other learning algorithms is the emergence of complex structures and behavior from strictly local interactions (Valiant, 2013; Watson and Szathmâry, 2016). The key molecular interactions responsible for faithful translation of RNA transcripts to proteins in contemporary living systems is the highly specific bindings of aminoacyl-tRNA synthetases (aaRSs) to their cognate tRNAs and amino acids (Chin, 2017). There are two classes of aaRSs that are distinct in domain organization and structure due to their unusual origins in two complementary strands of a single ancestral gene (Rodin et al., 2009; Rodin and Ohno, 1995). The twenty contemporary canonical amino acids distribute their specificity evenly and non-randomly among the two aaRS classes, with Class I amino acids being mostly non-polar and hydrophobic and Class II amino acids mostly polar and hydrophilic (Figure 5, A & B) (Carter et al., 2014; Nagel and Doolittle, 1995). Conservation of physio-chemistry within the two classes of amino acids is most parsimoniously explained by successive duplications of two ancestral aaRS genes, with the specificity of each new duplicated paralogous gene copy modified to bind a new but physio-chemically similar amino acid. Further, phylogenies of both aaRS classes are significantly auto-correlated with their cognate amino acids (Figure 5, A & B) (Nagel and Doolittle, 1995).

**Figure 5.**
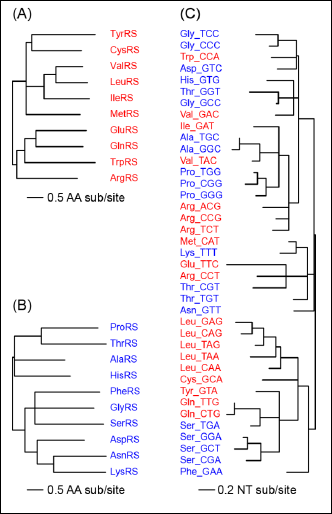
Molecular phylogenies of aaRSs and tRNAs from a *E.coli* genome. Molecular phylogenies of (A) Class I aminoacyl-tRNA synthetases (aaRSs) (in red), (B) Class II aaRSs (in blue), and (C) tRNAs (labeled with the cognate amino acid names and the anticodon sequences) from the *E.coli* K-12 genome. Only significantly supported branches are shown (see Methods). In Class I aaRSs, there is significant evidence for co-diversification between the aaRS sequences and the polarity of their cognate amino acids (Moran’s I = −0.044, p = 0.020 with n = 999 randomized samples). Test of phylogenetic autocorrelation for Class II aaRSs is similarly significant (Moran’s I = −0.075, p = 0.039). Evidence for co-diversification between rRNA sequences and cognate amino acids is weak for polarity (Moran’s I = 0.0077, p = 0.064) but strong for hydrophobicity (Moran’s I = 0.044, Ρ = 0.004). In sum, phylogenies of three key gene families involved in genetic code recognition all show clear footprints of primordial co-diversification between gene sequences and physio-chemistry of cognate amino acids. The correlated phylogenetic history between macromolecules and amino acids provides the biological basis for modeling the origin of SGC as a Hebbian learning process and is thereby called “phylogenetic learning”. Note a lack of apparent association between aaRS classes and tRNA clades, which suggests that while diversification of each gene is driven by recognition to amino acids these genes do not drive each other’s evolution.

#### tRNAs Phylogeny

Specific binding of aaRSs with their cognate amino acids accomplishes only half of the solution for faithful translation of codons into proteins. The other half - recognition of amino acids by codons - is mediated by tRNAs, each of which is ligated (by an aaRS) at its acceptor stem to a specific amino acid according to the anticodon sequence (Rodin and Rodin, 2008). Unlike the aaRSs, tRNAs are structurally homologous, suggesting an origin from a single ancestral coding sequence. Phylogenies of contemporary tRNA gene sequences show a general monophyly of iso-accepting tRNAs (i.e., tRNAs recognizing the same amino acids) although codon recruitments from different tRNA clades have occurred (Figure 5C) (Saks et al., 1998). Further, significant phylogenetic autocorrelation of tRNA with hydrophobicity of cognate amino acids supports that expansion of iso-accepting tRNA groups to all twenty amino acids involved similar tRNAs recognizing physio-chemically similar amino acids (Figure 5C).

Together, the phylogenetic associations observed in contemporary translation systems - the association between similar aaRS sequences and physio-chemically similar amino acids and the association of similar tRNAs with the same amino acids - are sufficient to confer an accurate mapping between 20 amino acids and 64 codons. The specific binding between the two macromolecules themselves (aaRSs and tRNAs) - the other pair of molecular recognitions - do not seem to have evolved through correlated phylogenetic co-diversification considering a lack of apparent association between aaRS classes and tRNA clades (Figure 5) (Nicholas and McClain, 1995).

## Discussion

This study is motivated by the proposition that algorithmic learning could lead to self-optimized, adaptive, and robust living systems even when natural selection is absent (Valiant, 2013; Watson et al., 2016). Using the origin of genetic code as an example, we show that it is indeed plausible for an error-minimizing genetic code to emerge through a Hebbian learning process without natural selection playing a role at system levels.

### TSP representations of genetic code

Our algorithm for finding optimal genetic codes consists of two main steps. In the first step, we formulate code optimization as a combinatory optimization problem of finding the shortest paths traversing the 20 amino acids and the translation stop signal. This Traveling Salesman Problem (TSP) interpretation of the genetic code lends itself to be solved with the Hopfield neural network (Bout and Miller, 1989; Hopfield and Tank, 1986). In the second step, we map the optimal tour positions emerged from the Hopfield network to a circular sequence of codon - a codon wheel (Figure 1) - that are based on two biological constraints. The first constraint is codon degeneracy for assigning 20 amino acids to 64 possible codons. The second constraint is a bias towards more transitions than transversions in DNA mutations. Note that although we construct the codon wheel by following these and other constraints present in SGC (e.g., the codon degeneracy decreasing in the order of 3^rd^, 1^st^, and 2^nd^ codon positions and the size distribution of synonymous codon blocks), there is no guarantee that optimized genetic codes emerged from the Hopfield network would be as optimal as SGC. This could be seen in randomly permuted codes, most of which have higher error rates than SGC although generated with the same set of constraints including codon-degeneracy, transition/trasnversion bias, and the size distribution of synonymous codons (Figure 2). More tellingly, plenty of codes optimized by the Hopfield network and mapped to codon wheel show higher error rates than SGC (Figure 3).

Together, these two algorithmic steps (one computational, one biological) allow us to establish that genetic code as optimal as SGC emerge quickly and non-selectively with Hebbian learning as the algorithmic principle operating within a set of biological constraints. It is conceivable that these two steps could be combined into a single TSP algorithm. We have represented the genetic code as a 64 amino acids (with repetition)-by-64 codons TSP and set the distances between the same amino acids as zero. In practice, however, optimal codes emerged from such a Hopfield network turned out to be not strictly comparable to SGC (and its random permutations) since the size distribution of synonymous codon blocks was not preserved. One way of preserving the codon block size distribution in SGC is to number the codon blocks sequentially according to a codon wheel (Figure 1) so that a 21 amino acid −21 codon block TSP could be constructed. This representation would be equivalent to the two-step algorithm we used since the “SGC address” of a codon is precisely the position of the codon block containing this codon in a codon wheel.

### Phylogeny recapitulates self-learning

The close analogy between an optimal genetic code and an optimal solution to TSP is not incidental, but due to the fact that both are manifestations of the Hebbian learning rule: an optimal tour of the cities is realized by traveling proximal cities in proximal tour positions, and an optimal genetic code is similarly realized by mapping similar amino acids to similar codons (Stoltzfus and Yampolsky, 2007). The Hebbian rule does not guarantee global optimum but does lead to locally stable shortest paths (Bout and Miller, 1989; Hopfield and Tank, 1986; MacKay, 2003).

Hebb’s rule enhances connections between neurons with correlated activities and weakens connections between uncorrelated neurons. Mathematically, the rate of increase in the weight (*w_ij_*) of connection between two neurons (*i* and *j*) with positively correlated (r) activities *x_i_* and *x_j_* could be expressed as: *dw_tj_/dt ~ r(x_it_X_j_*) (MacKay, 2003). This simple organization principle allows individual neurons to self-organize through auto-association, giving rise to a robust system capable of creating, storing, and retrieving a stable representation (“associative memory”) of an external pattern (e.g., an image) as shown by the example of a Hopfield neural network (Hopfield, 1982). Similarly, incremental co-diversification among aaRSs, tRNAs, and amino acids creates phylogenetic auto-correlations between gene sequences and amino acid physio-chemistry (Figure 5), giving rise to an effective and robust internal genetic representation (e.g., aaRS & tRNA sequences) of an external environment (e.g., amino acid physio-chemistry). In this sense, the gene phylogeny itself represent a molecular analog of the Hebb’s rule, which may be phrased simply as “similar paralogs bind similar amino acids”. Mathematically, the phylogeny-based learning rule could be expressed as: *dL_tj_/dt ~ d(AA_i_AA_j_*), where *L_ij_* is the phylogenetic distance (i.e., tree length) between two paralogs and *d(AA_i_, AA_j_*) is the difference (in e.g., polarity) between two amino acids they respectively bind. The stability of the protein translation system is maintained by the large and ever expanding phylogenetic distances between, e.g., Class I and Class II aaRSs or subclasses of tRNAs, creating an essentially digital and highly discriminating representation of amino acids that is SGC.

The Hebbian interpretation of the origin of SGC is consistent with the codon capture hypothesis, which posits that primordial codon expansion followed a phylogenetic path of minimum changes in amino acid physio-chemistry or exchangeability (Davis, 1999; Osawa and Jukes, 1989; Stoltzfus and Yampolsky, 2007). Extrapolating from the analysis of aaRS and tRNA gene phylogenies and the codon capture hypothesis, we propose that a gene phylogeny among paralogs represents a Hebbian learning process whenever a molecular system rises from co-diversification of gene sequences and gene functions. For such a system, while natural selection is critical for maintaining functions of individual genes (e.g., sub-functionalization with increasing discriminating power among aaRS and tRNA paralogs), it is not necessary that natural selection operates at system levels through competition among alternative systems.

### Broader implications

Molecular systems and pathways consisting of diversifying paralogs are not limited to genetic code and its supporting molecular machinery. In fact, genome and gene duplications coupled with subsequent neo-functionalization or sub-functionalization are a powerful, pervasive, and predominant mechanism of evolutionary innovation at genome levels (Graur, 2015; Ohno, 2013). Duplicative origin of α- and ß-hemoglobin genes in a vertebrate ancestor has led to the tetraheteromeric organization of the hemoglobin molecule conferring novel regulatory capacities for binding oxygen and carbon dioxide molecules (Graur, 2015). Duplication and subfunctionalization of homeobox genes, which encode transcription factors responsible for body patterning during development, have been associated with body plan diversification in bilaterian animals (Holland, 2015). Vertebrate olfactory sensing system consists of species-specific, rapidly evolving members of a gene family encoding odorant receptors that detect environmental chemicals in generalist, specialist, and combinatory fashions (Graur, 2015). Besides allelic polymorphisms, shifting gene expressions among a repertoire of duplicated surface antigen genes allow microbial pathogens such as *Trypanosome* and *Borrelia* to evade host innate and adaptive immunity (Norris, 2014; Taylor and Rudenko, 2006).

In each of these cases, a complex adaptive molecular system has evolved from incremental phylogenetic co-diversification between genes and gene functions. It is not surprising that systems directly involved in biotic and abiotic interactions consist of the largest and most rapidly evolving repertoire of duplication genes. The Hebbian interpretation of the origin of SGC and other molecular systems hints a general phylogeny-based learning algorithm for generating an efficient and accurate mapping between an organism’s genome and its living environment, without natural selection playing a significant role at system levels. Further, the Hebbian interpretation suggests computationally efficient ways in TSP and Hopfield networks to represent a molecular system digitally, predict its evolutionary trajectory, and improve its performance with a synthetic system.

## Declaration

### List of abbreviations

SGC: standard genetic code; Ti: nucleotide transition; Tv: nucleotide transversion; TSP: traveling salesman problem; aaRS: aminoacyl-tRNA synthetase; tRNA: transfer RNA

### Availability of data and codes

Source code is available as Open Source on Github at https://github.com/weigangq/code-by-tsp. including Perl/Python/R codes for simulated annealing, Perl codes for random permutation of genetic codes, and Perl codes for calculating fitness of a genetic code.

### Authors’ Contributions

OA, BT, and CD implemented Hopfield network algorithms, ran simulations, and produced graphs. WGQ conceived the project, performed statistical analysis, and drafted the manuscript.

## Acknowledgements

We thank Dr Lia Di for graphic assistance. We thank Dr Jeremy Draghi for introducing us to evolutionary learning and inspiring discussions. This work was supported by Public Health Service grants AI107955 (to WGQ) from the National Institute of Allergy and Infectious Diseases (NIAID) and the grant MD007599 (to Hunter College) from the National Institute on Minority Health and Health Disparities (NIMHD) of the National Institutes of Health (NIH) of the United States of America. The content of this manuscript is solely the responsibility of the authors and do not necessarily represent the official views of NIAID, NIMHD, or NIH.

